# Discover Novel RNA Targeting Small Molecules by Fluorescent Aptamer Screening

**DOI:** 10.64898/2026.06.23.734115

**Authors:** Ying Xu, Mengfei Du, Yanjiao Wang, Yi Xue, Hang Shi

## Abstract

Discovering small molecules targeting proteins represents a major effort in drug development. RNA, however, as a class of macromolecule that carrying out important regulatory roles in the cell as drug target, only received attention recently. Although several methods have been proposed, an easy to operate, fast and robust method is still lacking. We designed a generic florescence screening method by fusing the target RNA with a florescent aptamer (fusion RNA) and then carried out screening using high-throughput format (Fluorescent Aptamer Screening, FAS). In this work, we chose SL5 on SARS-Cov-2 5’UTR as the test target. SL5 is a conserved motif across several corona virus family members whose core is not prone to mutation. We screened 9528 compounds, successfully identified four molecules (Sertraline (hydrochloride), Samuraciclib (hydrochloride), Minocycline (hydrochloride), JG-98 bind direct to the full-length SL5 at micromolar or higher affinity. The design of FAS could be easily adapted to structured RNA motifs without prior knowledge of its 3D structural information. In addition, this work showed the possibility of developing generic drugs for RNA virus by targeting the conserved viral RNA genome and paved a new way for the discovery of small molecule drugs in combating human diseases.

## Introduction

RNA has been widely regarded as regulatory molecules other than being the messenger (mRNA) to transfer genetic information in transcription and translation. Its regulatory form includes microRNA (miRNA), small interfering RNA (siRNA), circular RNA (circRNA), long non-coding RNA (lncRNA), and untranslated region (UTR) etc. RNA also serves as the only genetic material to RNA viruses. The functional versatility of RNA is rooted in its diverse structure, innate flexibility and unique chemistry. To carry out its function, RNA may assume higher order structure and present characteristics surface to interact with other cellular factors. Like proteins, RNA mutations or malfunctions could give rise to human diseases (Qian et al. 2020). Traditional drug development aims to target proteins involved in the disease process, but some of them lack appropriate drug-binding pocket or catalytic activity, thus regarded as undruggable. In these situations, RNA provides an alternative target for future drug development and is currently receiving enormous attention. In addition, RNA viral genome, viral mRNA and bacterial RNA are also excellent targets for the development of new drugs.

Current RNA-targeting drugs can be divided into two classes: oligonucleotides and small molecules (Costales et al. 2020; Quemener and Galibert 2022). Oligonucleotide can base pair with its target RNA thus give high specificity. However, it comes with several disadvantages: it is difficult to administer, easy to degrade, and could induce immune responses, etc. On the contrary, small molecule drug is easier to deliver, easier to transport and store, produced at much lower cost, and with better stability. With mature technology in drug design, improvement and massive production, it showed unparalleled advantage. However, searching for RNA targeting small molecules significantly lagged behind with only a handful of antibiotics mostly target bacterial ribosomes known for decades (Drysdale et al. 2002). Not only until 2020, Risdiplam, the first FDA approved drug targeting SMN2 for the treatment of Spinal Muscular Atrophy (SMN) became available (Ratni et al. 2018).

Compared with the design of oligonucleotide which only requires targeting sequence, small molecules tend to bind structured regions of an RNA. Early work explored many simple motifs including internal loops, apical loops and bulges (Seth et al. 2005). However, these low complexity motifs typically yielded candidates with modest potency and selectivity. Complex, high-information-content structures such as multi-helix junctions and pseudoknots are proposed as better targets (Warner et al. 2018). In principle, if a 3D structure of an RNA is available, we could use computation tools developed for protein drug discovery to predict or even design binders (Aboul-ela 2010; Stelzer et al. 2011). Unfortunately, this is still difficult since only about 2000 RNA structures (as opposed to over 100,000 protein structures) are available with large number of them coming from ribosome or tRNA families, making it difficult to generalize. Moreover, the resolution of RNA structure tends to be low, giving ambiguous contact information. When we face a new RNA target, solving its 3D structure is time consuming, or a lot of time, unsuccessful due to the flexibility of the molecule. The bottle neck for discovering a drug for a new RNA target thus lies at the first step: how to quickly find candidates?

Both computational and experimental efforts have been made to discover candidates. Based on secondary structure obtained by SHAPE or 3D structure obtained by integrating SHAPE and modeling, people have used information obtained from known RNA binders (less than 300) to scan for candidate (Suresh et al. 2023). Given too few accurate structural data available, the effectiveness of virtual screen remains questionable. Developing efficient experimental assays may be the key to solve this problem. Although screening strategy based on the property of an individual RNA, for instance ribozyme (Fedorova et al. 2018) or frame shift activity (Sun et al. 2021), has been proven successful, tools that could be generically applied to variety of RNA remain very limited. Rizvi and Nickbarg developed the automated ligand identification system (ALIS) (Rizvi and Nickbarg 2019), in which they first incubate compounds with RNA followed by size-exclusion chromatography to purify any complex, then use liquid chromatography coupled to mass spectrometry to identify RNA binders. This procedure may give an advantage in finding ‘strong binders’ but leaving dynamic binder or ‘weak binder’ unidentified. Childs-Disney et al. developed the microarray assay in which compounds were linked to the gel plate followed by incubation with labeled RNA, washed and then read (Childs-Disney et al. 2018). This procedure requires RNA labeling, preparation of compound embedded plates, and washing steps, thus is technically demanding. More importantly, current available methods usually yielded candidates that primarily target bulges or loops that lack sequence and structural complexity, thus these small molecules very likely present low specificity in further analysis. To improve the specificity, drugs targeting complex structures such as the core of multi-way junction or pseudoknot are highly expected (Yu et al. 2020). Therefore, a method that could lead to the discovery of new candidates directly target the complex RNA and meanwhile take full advantage of high-throughput (HT) systems developed in protein drug research would provide its unique value.

In this work, we designed a fluorescent RNA aptamer (FRA) screen that could easily adapt to a desired RNA and subject to HT screen (Fig. 1). Fluorescent RNA adapters are a class of RNA (do not give off light) that could emit bright fluorescence when interacting with specific fluorophore (usually gives off very low signal). FRA has been widely used in detecting RNA localization in the cell, studying RNA folding, RNA-protein interaction, RNA riboswitch and designing small molecule biosensors etc (Ryckelynck 2020). In this study, we used Spinach 2 developed by Matthew D Disney & Samie R Jaffrey as a reporter (Strack et al. 2013). Spinach 2 is a 95-nucleotide-long RNA that could specifically interact with a GFP-like small molecule Fluorophore4-(3,5-difluoro-4-hydroxybenzylidene)-2-methyl-1-(2,2,2-trifluoroethyl)-1H-imidazole-5(4H)-one (DFHBI-1T) to emit bright green fluoresce at the excitation wavelength 482 nm and emission wavelength 505 nm. It contains a G-quadruplex (G4) at its core and a single stem-loop (SL). When its stem was shortened and properly fused to the stem of a structurally unstable target RNA to form a transducing stem, the folding of the G-quadruplex is disrupted so do its binding to DFHBI-1T, hence the emission of the Spinach 2 is attenuated. However, if a compound is bound to the target RNA which sufficiently stabilizes the transducing stem, Spinach 2 could be stabilized and enables the reading of interaction with an increased fluorescence signal. On the contrary, if the small molecule destabilizes the target RNA core, then the fluorescence signal is expected to reduce (Fig. 1). Thus, this method, when designed properly, allows the detection of candidates with both effects.

**Figure 1.**
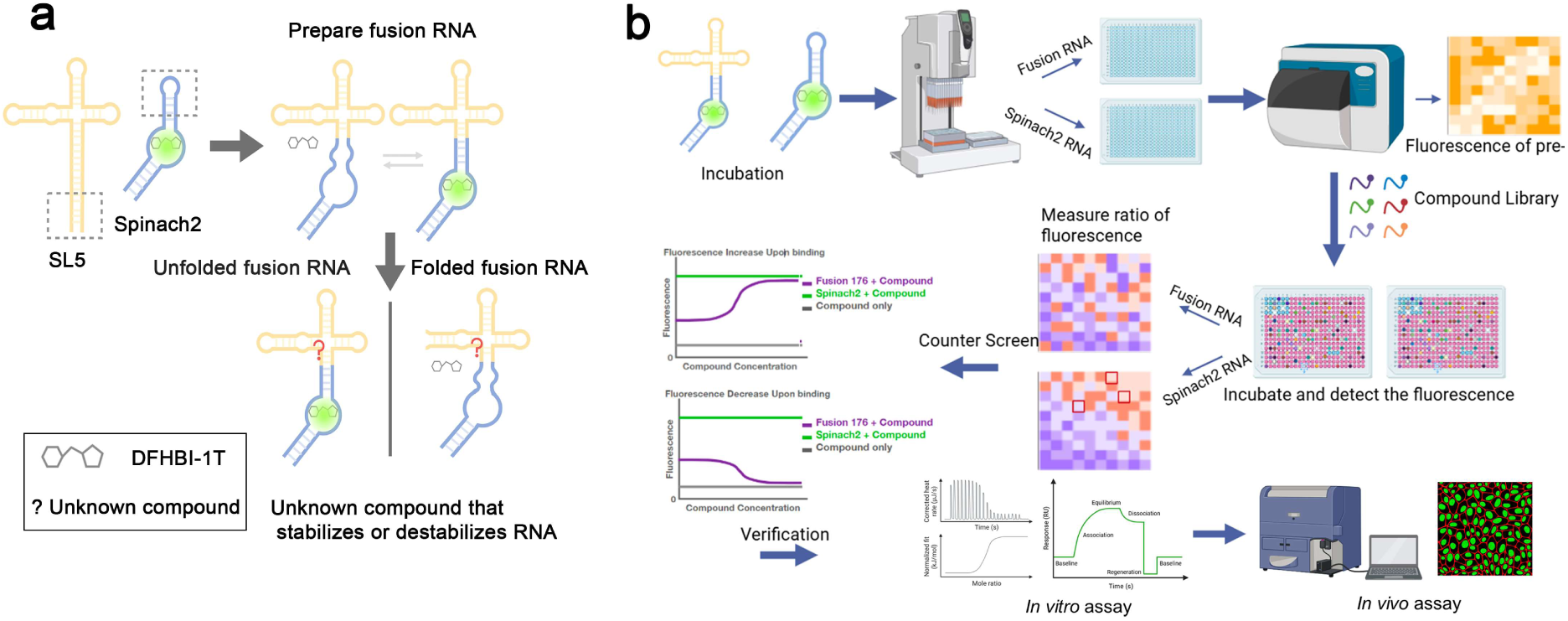
Schematic diagram of the design and process for small molecule screening using Spinach 2 and SL5. **(a**) Schematic diagram of the experimental design of FAS. The target RNA and the fluorescent RNA were connected through the transducing stem to form fusion RNA. The fusion RNA remained in dynamic equilibrium between the folded and partially folded states. When the fusion RNA was bound with a candidate small molecule, it could stabilize the structure of the target RNA, through which stabilized the transducing stem, thus moving the equilibrium toward the folded form and increased the fluorescence. Likewise, if the small molecule destabilized the target RNA, then the fluorescence would decrease. (**b**) The flowchart of screening and verification procedure. The fusion RNA was designed and prepared by *in vitro* transcription. The control group (Spinach 2 in this work) and the experimental group Fusion RNA were incubated and aliquoted into high-throughput plates, respectively. The basal fluorescence was measured by multi-functional microplate reader. After adding screening compounds, the fluorescence was measured again. Hit compounds were selected based on the change in fluorescence differences and ratios between the fusion RNA and the control. To further verify the efficacy of these compounds and eliminate false positives, the counter screen was first conducted to select top candidates. These candidates would be further verified against wildtype target RNA by *in vitro* binding assays such as ITC. Then the top candidates would be selected for subsequent experiments *in vivo* for functional verification.

In this work, we tested the SL5 motif from severe acute respiratory syndrome coronavirus 2 (SARS-CoV-2) 5’ UTR region (Hartenian et al. 2020) as the screen target. SARS-CoV-2 belongs to β coronavirus family that primarily infects mammals. It is the cause of COVID19 that lead to 13 million deaths during 2020-2023 pandemic (Llanes et al. 2020). Two other viruses in this family that also caused public health emergency in the past several decades were severe acute respiratory syndrome coronavirus (SARS-CoV,(Tratner 2003)) and middle east respiratory syndrome coronavirus (MERS-CoV, (Omrani et al. 2015)), highlighting the importance of finding generic drugs to combat this viral family. β coronaviruses are positive strand RNA viruses that contain a genome of about 30,000 nts (Llanes et al. 2020). At the ends of the genomes are short 5’ UTR and 3’ UTR crucial to the genome replication, packaging and viral proteins translation. In between are open reading frames that encode several proteins essential for its infection, replication, packaging and budding (Tan et al. 2012). Several small molecule drugs were developed for SARS-CoV-2 targeting two viral proteins: Nirmatrelvir/ritonavir (Hammond et al. 2024), Simnotrelvir/ritonavir (Ye et al. 2025), Ensitrelvir (Yotsuyanagi et al. 2024) and RAY1216 (Hu et al. 2025) inhibit 3-chymotrypsin like protease (3CLpro), the main protease that cleaves the polyproteins into functional units; Remdesivir (Bakheit et al. 2023) and Molnupiravir (Syed 2022) inhibit RNA-dependent RNA polymerase (RdRp) to prevent viral replication. β coronavirus family genome is highly prone to mutations (Chen et al. 2021), to face future thread, a new direction for drug search is to target highly conserved regions in viral RNA genome to acquire wide spectrum drugs for β coronavirus family. For instance, Matthew D. Disney et al. found small molecules to target frameshifting element (FSE) in SARS-CoV-2 genome which inhibit proper translation of viral proteins (Haniff et al. 2020a). In addition, Ben F. Luisi at al. designed (anti-sense oligonucleic acid) ASO aiming for SL structures in 3’ UTR (Lulla et al. 2021). Researchers from Alnylam Pharmaceuticals designed 350 siRNA and try to identify effective ones that could target viral genome (Egli and Manoharan 2023).

The 5’ UTR of SARS-CoV-2 is highly structured and carries important functions (Santos-López et al. 2021). It forms five SL structures namely SL1-SL5. Each SL hosts a unique function, among them SL5 (nt 149-296) represents the most complex one (Fig. 2a and b), (Chen et al. 2021). Crystallographic and cryo-electron microscopy analyses reveal that SL5 adopts a dynamic T-shaped four-way junction (4wj) configuration, in which multiple helical segments converge to form a compact yet flexible three-dimensional fold. This junctional architecture creates well-defined deep pockets and grooves, arising from the spatial arrangement of helices and loop regions. These pockets are structurally pre-organized yet exhibit conformational adaptability, reflecting the intrinsic mobility of the SL5 RNA. Such features are particularly noteworthy, as they provide potential binding sites for small molecules, enabling specific recognition through shape complementarity and local chemical interactions. The combination of structural stability, conserved architecture, and the presence of ligand-accessible pockets highlights SL5 as a promising RNA target for structure-based drug discovery(de Moura et al. 2024; Kretsch et al. 2024; Jones and Ferré-D’Amaré 2025). Three SL extending from the major stem are named SL5a-SL5c, respectively (Fig. 2a and c). Two UUYYGY loops reside at the tips of two SLs (SL5a and SL5c), respectively, with 3.3 Å apart (Fig. 2a and c, (Jones and Ferré-D’Amaré 2025)). Importantly, the start codon of the first viral ORF is embedded in the SL5 major stem (Fig. 2a), suggesting its role in translational regulation. Indeed, when substructure of SL5 was deleted, the translation of genomic RNA (gRNA) would increase (Leppek et al. 2022). In the translation of the first ORF (ORF1a or ORF1ab), virus utilizes host translation system and at the initiation step. The 40S ribosome subunit recruits initiation factors to form 43S initiation complex and scans the 5’ UTR until it reaches the first ATG. In this process, the structure of SL5 is melted, allowing its association with 43S. The binding efficiency of initiation complex to the start codon is limited by the melting free energy and when a ligand is present to increase the melting free energy, the association rate to the 43S is reduced, thus inhibit the translation (Ray et al. 2022). In addition, SARS-Cov-2 SL5 can interact with N protein, a protein essential in viral packaging, suggesting it may also play a role in viral packing (Escors et al. 2003; Korn et al. 2023). Motifs like SL5 are also found in SARS-CoV, MERS-CoV viruses (Fig. 2a and b, (Chen et al. 2021)). Furthermore, the sequence of SL5 is nearly identical to SARS-CoV and its 4WJ structure is among the most conserved features of β coronavirus family (Fig. 2a and b, (Llanes et al. 2020)). Although the RNA genome of SARS-CoV-2 is highly prone to mutations, only a few were observed on SL5 (Saldivar-Espinoza et al. 2023), mostly at the distal ends of substructures (Fig. 2a), emphasizing its functional importance. Therefore, it serves an ideal target for generic drug development.

**Figure 2.**
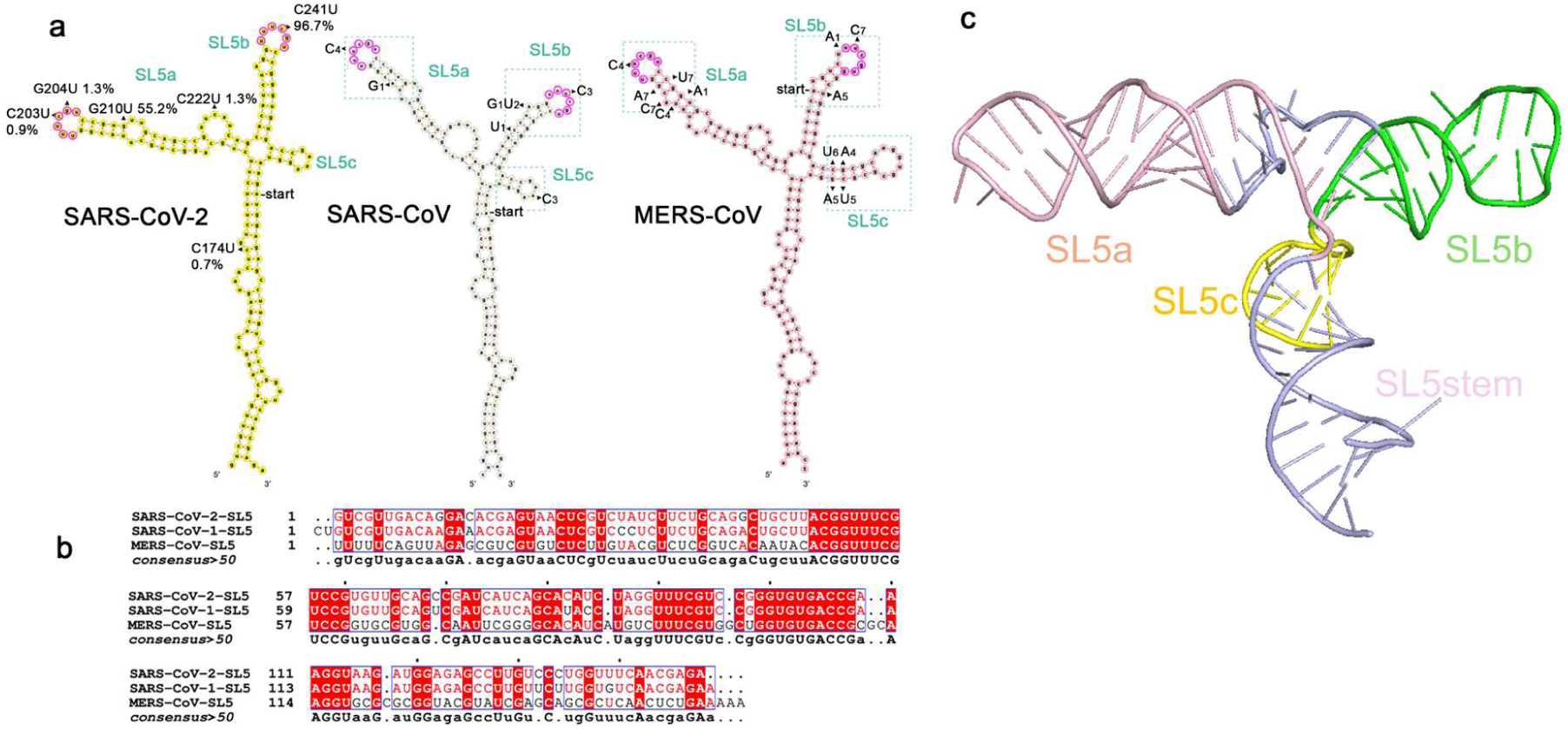
Sequence and structural conservation of SL5 in SARS-CoV, MERS-CoV, and SARS-CoV-2. **(a**) The secondary structure and single nucleotide variations of SL5 in three representative coronaviruses. The UUYYGU conserved sequence at the apical regions of SL5a and SL5b were marked by purple and dashed boxes. The secondary structure was drawn based on the cryoEM structures (PDB code: 8UYS (Kretsch et al. 2024)). The single nucleotide variation type and frequency were indicated and marked by triangle. The type and frequence for SARS-CoV-2 were retrieved from the SARS-CoV-2 Mutation Portal website (Saldivar-Espinoza et al. 2023), and that of SARS-CoV and MERS-CoV were reported previously (Chen et al. 2021). (**b)** Sequence conservation of SARS-CoV, MERS-CoV, and SARS-CoV-2. Sequences with 100% identity were colored with red background and letters in red indicated over 50% sequence identity. (**c**) Three-dimensional structures of SL5 from SARS-CoV-2 (PDB code 9E9Q,(Jones and Ferré-D’Amaré 2025)).

In this work, we applied FAS on SL5 of SARS-CoV-2 and identified four molecules out of 9528 as potential binders. Isothermal Titration Calorimetry (ITC) analysis confirmed their direct interaction to full-length SL5 with high affinity. Three of them also showed interaction with SL5 from SARS-COV and MERS-CoV, supporting the initial goal of this strategy to identify potential generic drugs.

## Results

### Design of SL5 fusion RNA

In this study, the classic fluorescent RNA Spinach 2 was chosen to produce the fusion RNA, with DFHBI-1T as its corresponding fluorophore (Strack et al. 2013). The Spinach 2–DFHBI-1T complex exhibits a fluorescence spectrum and intensity similar to those of green fluorescent protein, providing sufficient brightness and a suitable dynamic range for fluorescence measurements on instruments (Song et al. 2014).

We designed the fusion RNA of SL5 by truncating the basal stem of the SL5 below the junction by more than 30 base pairs and fused it with Spinach 2 SL with truncation from 36 to 58 nts (Fig. 3a). The varying truncation positions resulted in fusion RNA with different transducing stem length, hence giving off different brightness. To determine the ideal screen target, a total of eight fusion RNAs with different stem lengths were tested. We named the fusion RNA, formed by truncating at position 176 in the 5’ UTR, as Fusion 176, which contained an SL5 length of 94 nts, and similarly, designed and synthesized Fusion 171, 175, 177, 178, 179, 180, and 182 with an SL5 length of 99 nts, 95nts, 93nts, 92nts, 91nts, 90nts and 88nts respectively (Supplementary Fig. S1). The transducing stems of these constructs vary in length of 11, 8, 6, 5, 4, 3 and 1 base pairs.

**Figure 3.**
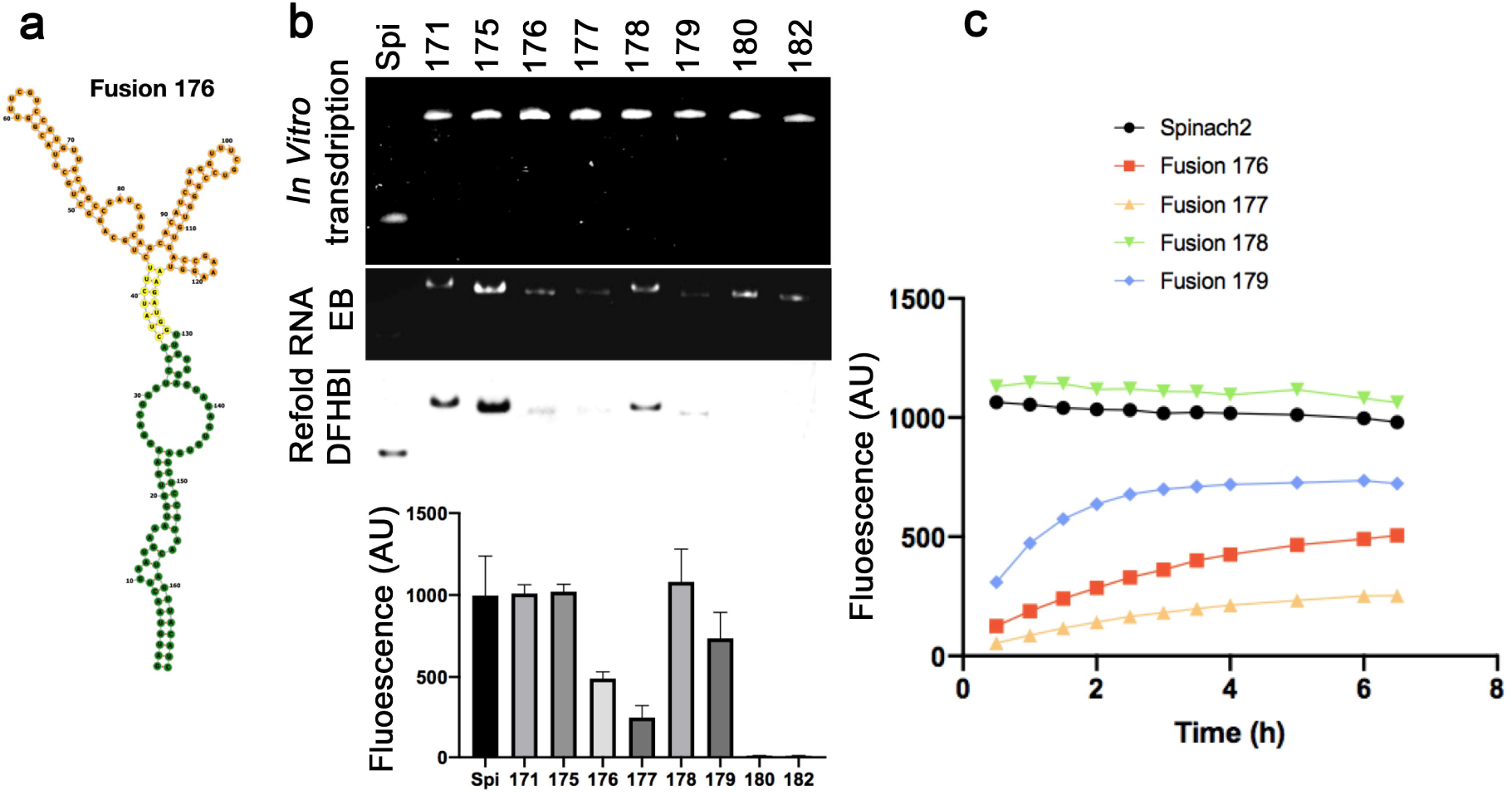
The design of Fusion RNA and the determination of screening conditions. **(a**) The design of SL5 fusion RNA for FAS. The SL5 of SARS-CoV-2 with truncation at the basal stem starting from position 176 was fused to Spinach 2 with truncation from 36 to 58 nts to produce Fusion 176. The yellow, green, and orange indicate the transducing stem, Spinach 2 and the region of SL5 excluding the basal stem, respectively. (**b)** Electrophoresis of Spinach 2 and fusion RNA. Spinach 2 and fusion RNA were subjected to PAGE electrophoresis and stained with ethidium bromide and DFHBI-1T, respectively. Spi represented Spinach 2, whereas 171、 175、176、178、180、182 represented fusion RNA with the corresponding truncation position at the basal stem. c) Pre-incubation was required to stabilize the fluorescence. Various incubation time (from one to seven hours) for Spinach 2 (black), Fusion176 (coral), Fusion177 (orange), Fusion178 (green), Fusion179 (blue) was required for the fluorescence to reach the maximum.

### Optimization of the screening condition

Previous studies have established preferred conditions for fluorescence measurements of the Spinach 2–DFHBI-1T complex, including buffer preparation and the concentrations of RNA and DFHBI-1T. The fluorescence detection conditions for the fusion RNAs in this study were based on these earlier protocols with further validation. We tested fluorescence at RNA concentrations ranging from 30 nM to 1 µM and DFHBI-1T concentrations from 1.7 to 20 µM (Supplementary Fig. S2). Additionally, we evaluated fluorescence at Mg²⁺ concentrations from 0 to 17.3 mM (Supplementary Fig. S2), considering that Mg²⁺ may influence RNA folding (Strack et al. 2013; Manna et al. 2021). In addition to RNA, DFHBI-1T, and Mg²⁺, the reaction mixture also contained 40 mM HEPES and 125 mM K⁺ in the imaging system. HEPES provides stable pH buffering for RNA, while K⁺, as the primary intracellular cation, creates a stable ionic environment for RNA (Manna et al. 2021).

Consistent with previous studies, RNA concentration exhibited a linear relationship with fluorescence intensity within the tested range (Supplementary Fig. S2a). In contrast, DFHBI-1T concentration did not affect fluorescence (Supplementary Fig. S2b). These results indicated that DFHBI-1T was saturated in the tested concentration range while RNA was not. Furthermore, Mg²⁺ concentration had no significant effect on fluorescence levels (Supplementary Fig. S2c), suggesting that Mg²⁺ has minimal impact on the folding of the RNAs tested. Finally, we determined the optimal concentrations for the system containing 300 nM RNA, 3 µM DFHBI-1T, and 10 mM Mg²⁺. Under these conditions, DFHBI-1T was saturated, and the relatively high RNA concentration facilitated a more sensitive response to changes in RNA folding states. Other than these, the buffer also contains 40 mM HEPES-KOH pH 7.5 and 125 mM KCl.

We later measured the fluorescence values of fusion RNAs (Fig.3). The longest fusion RNAs, Fusion 171 and Fusion 175, exhibited fluorescence levels comparable to Spinach 2, while the shortest fusion RNAs, Fusion 182 and Fusion 180, showed almost no fluorescence (Fig.3). This aligned with the prediction that longer transducing stem stabilized the RNA thus enhancing the fluorescence. Fusion 176, Fusion 177, Fusion 178, and Fusion 179, which had intermediate transducing stem lengths, displayed varying fluorescence intensities (Fig.3).

We further examined the series of fusion RNAs using non-denaturing PAGE electrophoresis, staining with either ethidium bromide (EB) or DFHBI-1T (Fig. 3b, middle and bottom panel). The brightness of each RNA in the DFHBI-1T-stained gel agreed with the fluorescence measurements obtained using a plate reader. From Fusion 171 to Fusion 182, the migration speed of the RNA on the gel progressively increased as the RNA sequence and transducing stem length decreased. The sole exception was Fusion 178, which migrated slower than other fusion RNAs of similar length. However, denaturing PAGE confirmed that the migration rate of Fusion 178 was normal (Fig. 3b, top panel). This suggested that Fusion 178 may possess a unique tertiary structure, resulting in its distinct migration speed and relatively high fluorescence level. Based on these results, we selected Fusion 176 as our screening target to search for compounds that could stabilize or destabilize the SL5 structure.

During fluorescence testing, we observed that the incubation time of RNA with DFHBI-1T also influenced the read out (Fig. 3c). Previous studies have reported that Spinach 2 (indicated as Spi in Fig. 3) reached stable fluorescence within one hour of incubation with DFHBI-1T (Anisuzzaman et al. 2022), and our findings were consistent with this conclusion. For fusion RNAs with fluorescence levels comparable to Spinach 2, such as Fusion 178, fluorescence also stabilized within one hour. However, for fusion RNAs with lower emissions, fluorescence gradually increased over several hours until stabilization (Fig. 3c). This phenomenon may reflect a process in which fusion RNAs were partially folded and the DFHBI-1T binding to the correctly folded RNA shifted the folding equilibrium, driving more RNA molecules to adopt the correct conformation. The time required for fluorescence stabilization thus likely reflects the time needed for the refolding of the fusion RNA.

### Initial Screening of SL5 and counter screens

In this work, the MCE Bioactive Compound Library of the Active Screening Platform was used for screening, and a total of 9,528 compounds were screened. The control group (Spinach 2 in this work) and the experimental group Fusion 176 were incubated and aliquoted into high-throughput plates, respectively. The basal fluorescence was measured by multi-functional microplate reader. After adding screening compounds, the fluorescence was measured again. When the compound exhibited significant fluorescence, we selected initial hit compounds based on the changes in fluorescence differences that is, the fluorescence intensity after adding the compound minus the intensity before the addition; and when the compound showed no inherent fluorescence, we selected by the ratio of relative fluorescence changes that is, the ratio of the fluorescence intensity after the addition to that before the addition. The combination of these two calculation methods can comprehensively analyze the influence of the compound, eliminate errors caused by the non-specific binding of the compound to Spinach 2 or the inherent fluorescence of the compound. We initially identified 190 possible compounds, among which 67 increased the fluorescence of Fusion 176 compared to Spinach 2 alone (Fig. 4a and b), another 123 compounds showed reduction in fluorescence with the extent of the reduction in Fusion 176 different from that in the Spinach 2 (Fig. 4a and b). To further verify the efficacy of these compounds and eliminate false positives, we conducted counter screen verification.

**Figure 4.**
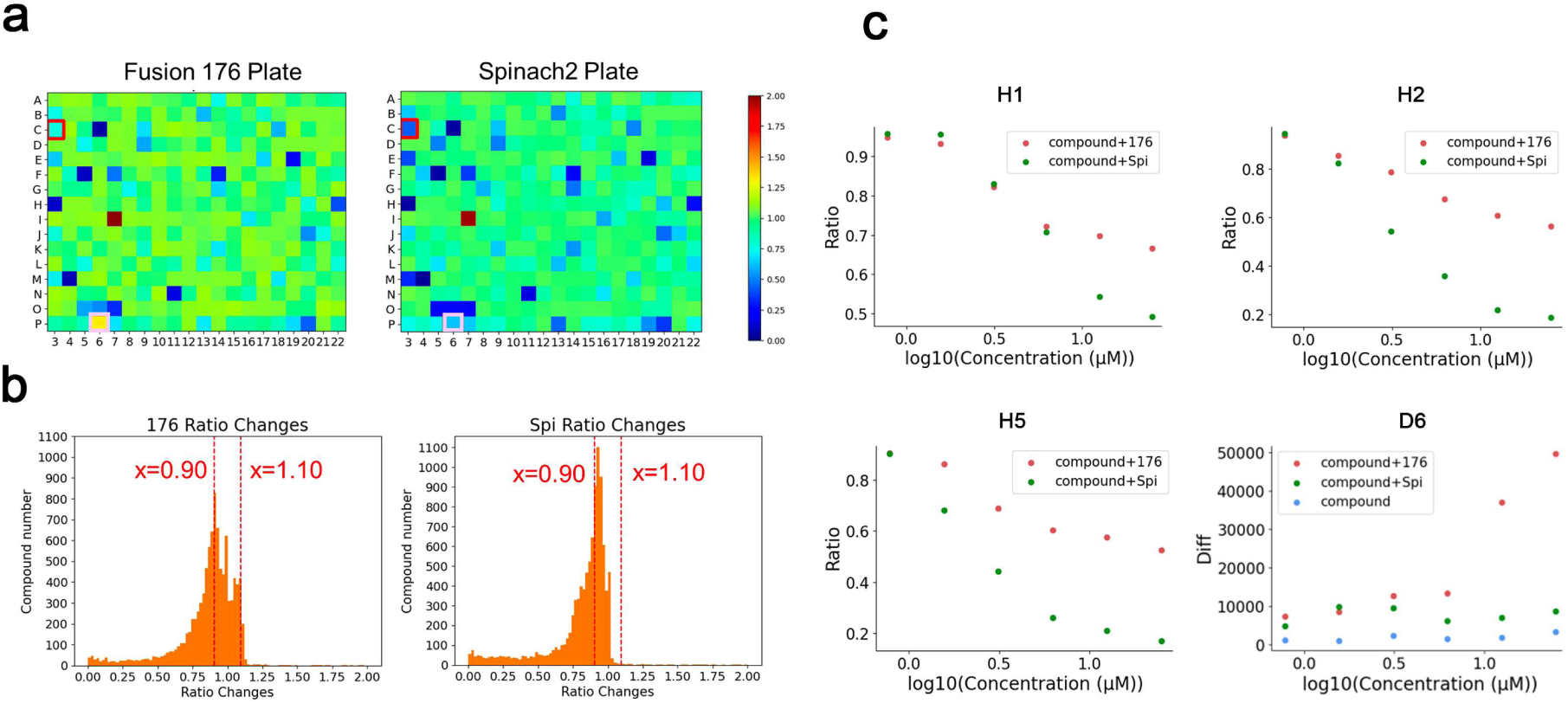
The hit compounds selected by FAS. **(a**) A representative illustration of the fluorescence difference between Spinach 2 and Fusion 176. Different compounds were added into each well. The color indicated the ratio of the fluorescence intensity between after addition of the compound and before the addition. The color bar to the right corresponded to the variation of the ratio. The representing plate showed the fluorescence increase (pink box) or decrease (red box) between Fusion 176 and Spinach 2. (**b)** The statistics of fluorescence changes of all screened compounds. The fluorescence responses of the Fusion 176 and Spinach 2 to all the screened compounds were plotted with x-axis indicated ratio change of the fluorescence intensity and y-axis indicated number of compounds, respectively. The red dotted lines indicated the positions of approximately two-fold of the CV with x=0.90 (left) and x=1.10 (right), respectively. (**c)** The results of counter screen of hit compounds. H1, H2, H5 led to the decrease of fluorescence in both Fusion 176 (red line) and Spinach 2 (green line). The difference between Fusion 176 and Spinach 2 dramatically increased following the increased compound concentration. The x-axis indicated the compound concentration whereas the y-axis indicated the ratio change of the fluorescence intensity. The D6 (with its own fluorescence, the blue line) caused a significant increase in fluorescence in Fusion 176 groups (red line), while there was no significant change in the Spinach 2 group (green line). The x-axis indicated the compound concentration whereas the y-axis indicated the difference change of the fluorescence intensity.

In counter screen, gradients of six compound concentration were set from 0.78 µM to 25 µM, and a control group without RNA was also set up to eliminate the influence of the compound’s fluorescence on the experimental results. Among the 67 compounds that led to increased fluorescence, only one compound (D6) caused a significant increase in the fluorescence of the Fusion 176 group, while no significant change in the Spinach 2 was observed (Fig. 4c). The remaining compounds did not cause significant differences in fluorescence between Fusion 176 and the Spinach 2.

Among the 123 compounds that led to fluorescence decline, 49 compounds demonstrated a smaller fluorescence decline in the Fusion 176 than that in the Spinach 2, and this gap widened with the increase in compound concentration (Fig. 4c, e.g. H1, H2, H5). This phenomenon may have two explanations. In one scenario, the SL5 structure in the Fusion 176 had a specific response to the compound, which was reflected in the difference of the fluorescence decline ratio. This compound was a potential hit compound that could stabilize the SL5 structure. In the other scenario, all these compounds could disrupt the fluorescence of Spinach 2. It was possible that the degree of their influence on Spinach 2 in the Fusion 176 was different from that on Spinach 2 alone, resulting in different levels of decline. To clarify whether these compounds directly interacted with SL5, we further conducted ITC to directly detect the binding of SL5 to the compound. The remaining 74 compounds achieved the same decline level between the Spinach 2 and the Fusion 176 in counter screening verification.

### Isothermal Titration Calorimetry

From the 49 compounds above that showed a reduced fluorescence decline, we used two criteria to select the top 10 compounds for ITC analysis (Supplementary Fig. S3c): firstly (and primarily), △Emax (difference in maximum response value), defined as the difference between the effect of the compound on Fusion 176 and that of Spinach 2 at the highest concentration; secondly, the IC50, to show the true effect of a compound by subtracting its effect on Spinach 2 from that on Fusion 176 (Supplementary Fig. S3a and S3b). We set the threshold at △Emax≥0.2 and IC50≤4.5. In addition, we also examined the one fluorescent compound that showed significant fluorescence increase between the Fusion 176 and Spinach 2.

Among 11 selected compounds, 10 compounds confirmed direct interaction with SL5, in which four compounds showed the combined K_d_ above 10 µM and the stoichiometry close to 1:1. These four compounds were Sertraline hydrochloride (H1, K_d_=5.6 μM), Samuraciclib hydrochloride (H2, K_d_=6.55 μM), Minocycline hydrochloride (H5, K_d_=669 nM) and JG-98 (D6, K_d_=7.3 μM), respectively (Fig. 5).

**Figure 5.**
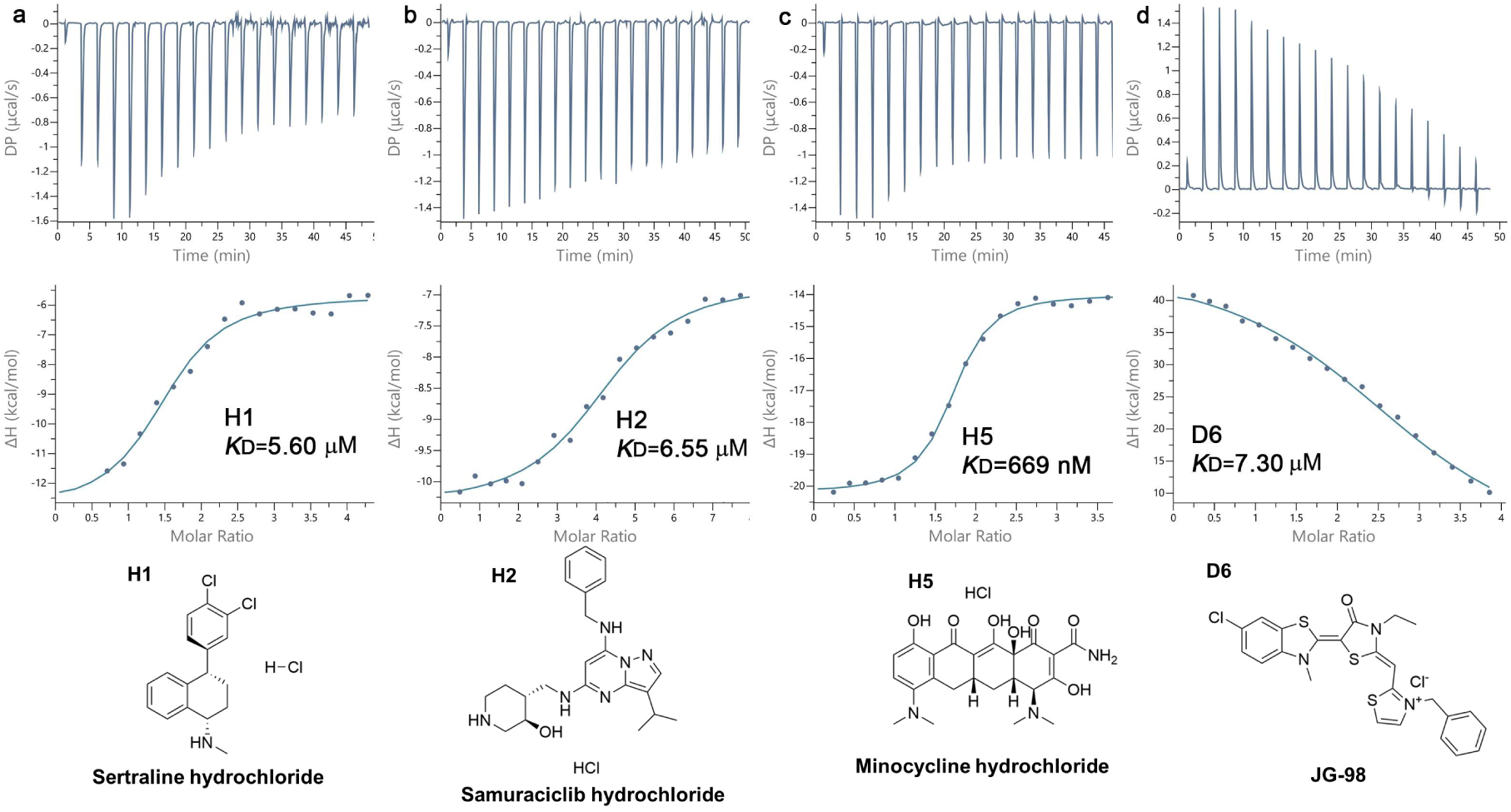
H1, H2, H5 and D6 bind the full-length SARS-CoV-2 SL5 *in vitro.* The Interaction between SARS-CoV-2 SL5 and H1 (left panel), H2 (middle left panel), H5 (middle right panel), and D6 (right panel) were significant. The thermodynamic titration curve was shown on top with the time plotted as x-axis and DP (Differential Power) for heat change plotted as y-axis, and the binding curve was shown at the bottom with compound concentration plotted as x-axis and enthalpy change plotted as y-axis. All four measured K_d_ were indicated with the corresponding binding curve. Each experiment was repeated three times. The chemical structures of H1, H2, H5 and D6. Structures of H1, H2, H5 and D6 were plotted by ChemDraw (Evans 2014)

### Compounds bind SL5 near the core of the 4wj

After we identified four candidate compounds that bind to SL5 directly, we next explored the specificity of the binding to obtain information on the binding sites by constructing two deletion mutants. Based on the reported structure of SL5 (Kretsch et al. 2024), we deleted several base pairs in SL5a and SL5b to obtain two mutants DA and DB (Fig. 6a and b, left panels), respectively. H1, H2, H5 could still bind to these two mutants, although the binding affinity was slightly lower than that with SL5, indicating that these deletions did not abolish the binding sites, and the compound may interact with SL5 within or near the 4wj core. A possible explanation to the reduced affinity may arise from the destabilization of the core which prevented the drug from optimal binding.

**Figure 6.**
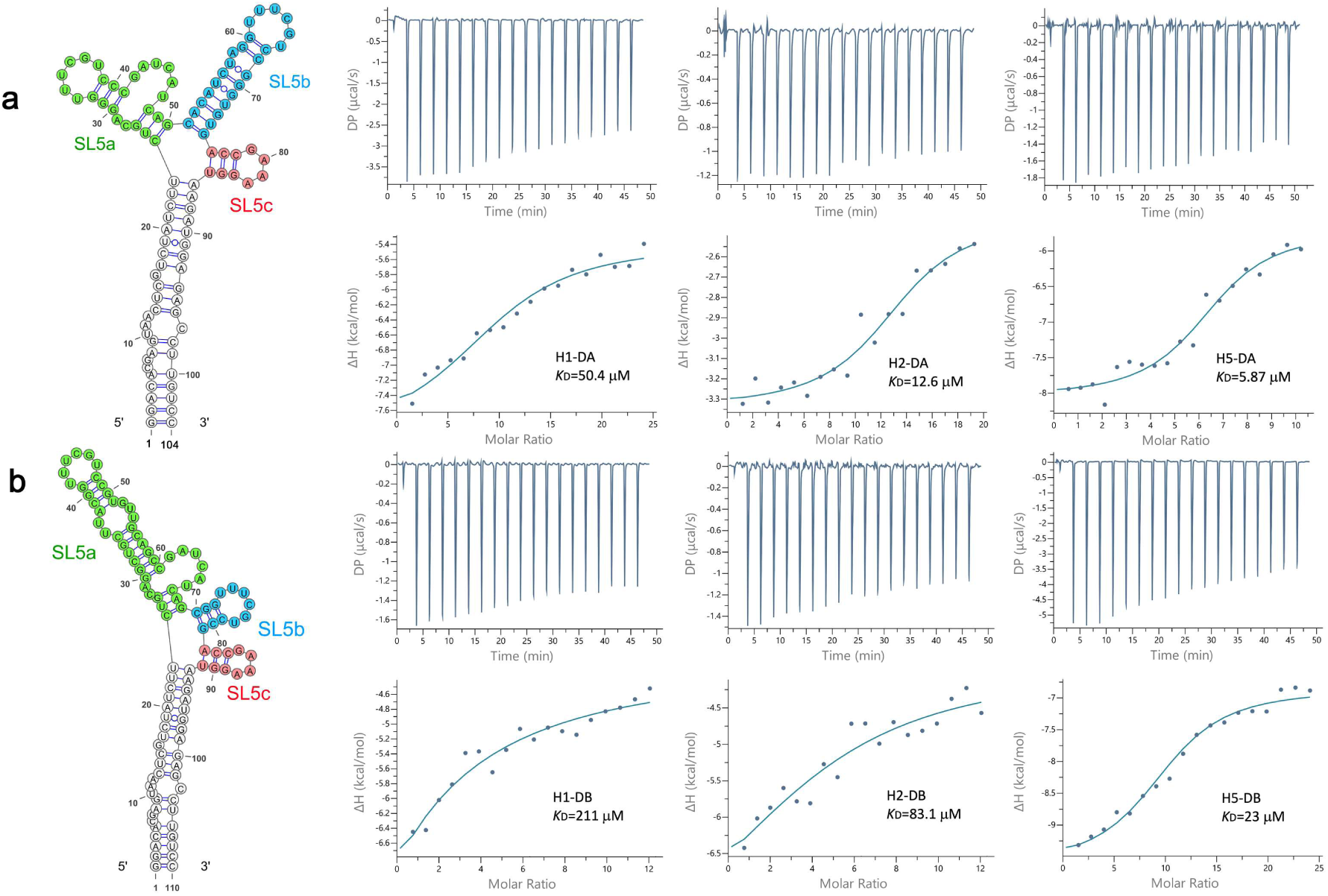
The deletion at SL5a or SL5b of SL5 reduced its interaction with H1, H2 and H5. **(a**) Deletion at SL5a reduced the interaction between SL5 and H1, H2, and H5. The secondary structure of SL5 with deletion at SL5a (DA) was shown at the left. The Interaction between DA and H1 (middle left panel), H2 (middle right panel), and H5 (right panel) were reduced when compared with that of the full-length SL5. The thermodynamic titration curve was shown on top with the time plotted as x-axis and DP (Differential Power) for heat change plotted as y-axis, and the binding curve was shown at the bottom with compound concentration plotted as x-axis and enthalpy change plotted as y-axis. All three measured K_d_ were indicated with the corresponding binding curve. Each experiment was repeated twice. (**b)** Deletion at SL5b reduced the interaction between SL5 and H1, H2, and H5 significantly. The secondary structure of SL5 with deletion at SL5b (DB) was shown at the left. The Interaction between DB and H1 (middle left panel), H2 (middle right panel), and H5 (right panel) were significantly reduced when compared with that of the full-length SL5. The thermodynamic titration curve, the binding curve and the K_d_ corresponding to each compound were shown as in **a**.

To further identify the binding sites, we performed ^19^F NMR titration experiments. SL5 and DA were selected as targets. Four ^19^F probes were simultaneously introduced at distinct positions within the junction regions of SL5 and DA (denoted as SL5-junction and DA-junction, supplementary Fig. S8). In addition, three ^19^F probes were placed at sites distal to the junction in DA as a negative control (denoted as DA-negative, Fig S8). The measurement relies on the fact that small-molecule binding near a ^19^F probe induces peak intensity reduction, chemical shift perturbation, or both, depending on *k*ₒₙ and *k*_off_.

As shown in Fig. 7, no significant changes in ^19^F spectra were observed for H1 and H2 upon binding to DA-negative, indicating no interaction. In contrast, for SL5-junction and DA-junction, both peak shifts and pronounced intensity decreases were observed, confirming binding. Upon addition of H5, precipitation was observed for all three RNA constructs, along with a decrease in ^19^F peak intensity. No chemical shift changes were detected for DA-negative, whereas clear shifts were observed for SL5-junction and DA-junction (particularly pronounced for the latter). These results indicate that H5 does not bind to DA-negative but binds strongly to the other two constructs. The observed precipitation is most likely caused by interactions between H5 and the junction region, leading to reduced peak intensities across all constructs. In summary, ^19^F NMR demonstrated that H1, H2, and H5 bind to SL5 and DA, with binding occurring at or near the junction region, and that H5 exhibited stronger binding than the other two compounds.

**Figure 7.**
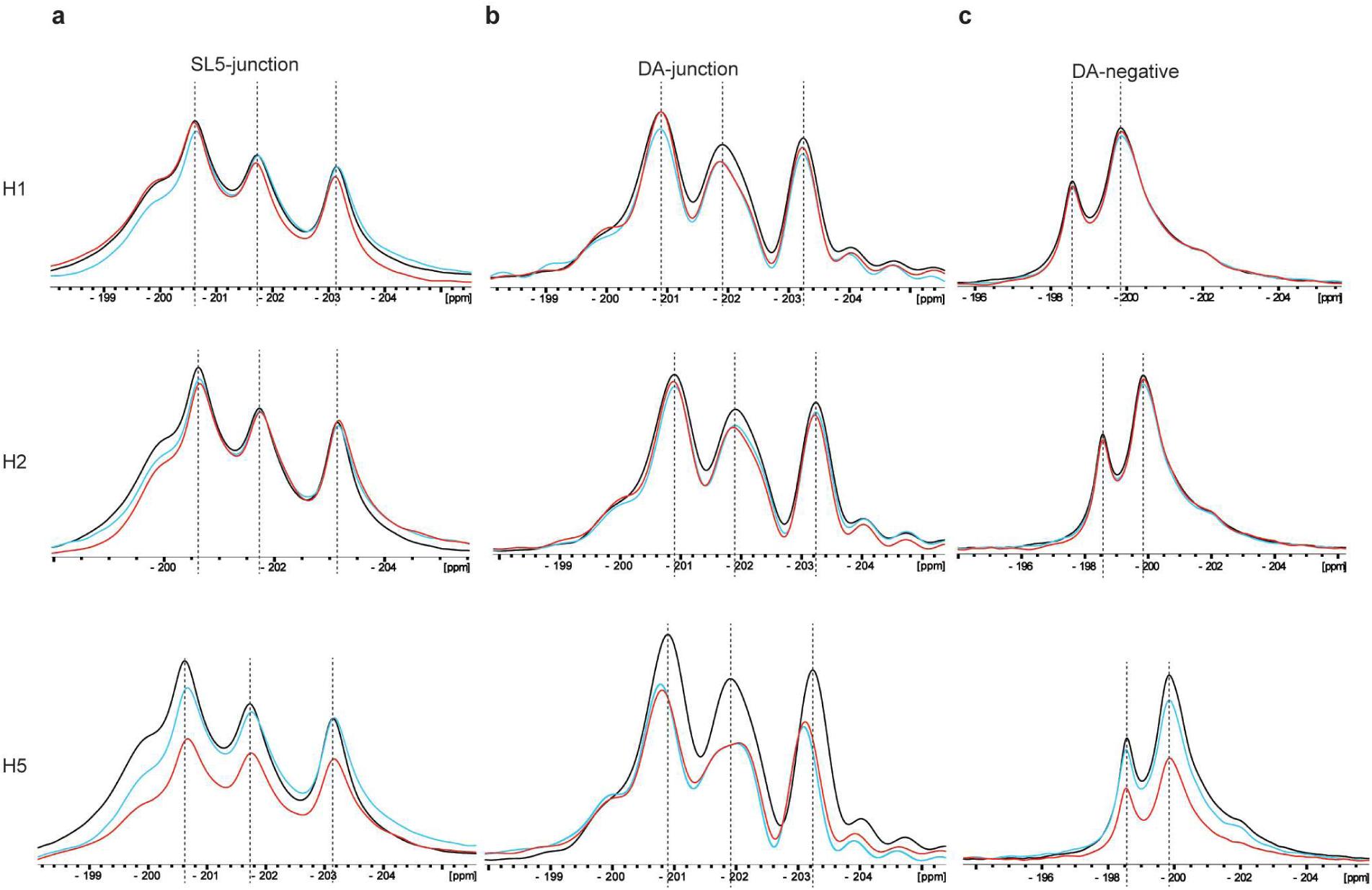
SL5 NMR titration experiments with H1, H2 and H5. NMR titration experiments for SL5-junction (a), DA-junction (b) and DA-negative (c) with H1, H2 and H5. SL5-junction and DA-junction were 2’F labeled at residue 222, 227, 257 and 262. DA-negative was 2’F labeled at residue 200, 233 and 242 (Fig. S8). The 1D spectra for RNAs titrated with small molecules at ratio 1:0, 1:1 and 1:3 were shown in black, blue and red, respectively.

To see whether H1, H2 and H5 interact with SL5 specifically, we conducted the ITC experiment using these three compounds to titrate bulk yeast tRNA under the same condition. The experiment did not reveal the binding of small molecules to tRNA (Supplementary Fig. S4), which confirmed that they did not widely bind RNA through non-specific forces such as electrostatic interaction or base accumulation, but selectively recognized a certain structure or sequence of the target molecule. Unlike the three compound above, D6 did not bind to either DA (Supplementary Fig S5a top panel) or DB (Supplementary Fig S5a bottom panel), indicating that its binding to SL5 may require a specific local sequence or structure.

### Three compounds interact directly with SL5 from SARS-CoV and MERS-CoV

The structures of SL5 in SARS-CoV-2, SARS-CoV and MERS-CoV are highly similar (Fig. 2a, (Chen et al. 2021)). Therefore, we further studied whether these four hit compounds could bind to SARS-CoV and MERS-CoV, that is, whether these small molecule drugs might have broad antiviral effects. Indeed, H1, H2, and H5 could all bind to SL5 of SARS-CoV and MERS-CoV, although the affinities decreased somewhat (Supplementary Fig. S6). These indicated that these three compounds have certain selectivity in binding to SL5, but the binding mechanism is sensitive to local structural changes. The binding of small molecules to RNA depends on certain fine conformations or hydrogen bond networks. Due to the slightly different tertiary structures or local conformations of these three viral motifs (Rabaan et al. 2020), it is very likely that the entry of small molecules into the corresponding sites has been affected. The complete non-binding of D6 (Supplementary Fig S5b) indicated that its binding is highly specific, and the subtle differences in conserved regions among different viruses may be sufficient to abolish the interaction.

### SL5 increases the fluorescence of D6

Surprisingly, the fluorescence of the D6 increased significantly after binding with Fusion RNA while no significant change was seen with control Spinach 2 (Fig. 4c). This suggested that the fluorescence of the compound may increase after binding with SL5. To test this, we compared the fluorescence of D6 in the presence of wildtype SL5 or in the absence of RNA. The results indicated that when combined with SL5, the fluorescence intensity of D6 could increase by more than two-fold at saturation (Supplementary Fig. S7a). Our further fluorescence scanning showed that the combination of SL5 and D6 didn’t alter the excitation and emission wavelengths of D6 (Supplementary Fig. S7b and c).

## Discussion

FAS is developed for structured RNA. A minimal stem loop is needed to use this system. However, FAS showed an incomparable merit on RNA with complex structure (Haniff et al. 2020b). In our work, we designed the transducing stem near the 4wj of SL5, a conserved feature also found in other corona viruses such as SARS and MERS. Upon drug binding, a sufficient conformational change that stabilized the transducing stem was pre-requisite to observe the florescent signal change. This represented an advantage in drug searching in that these molecules are more likely to bind at or near the designated structural features, hence possibly yielding higher specificity. Depending on the design, however, one drawback of this method was that it may miss binding sites that could not induce significant conformational change at the transducing stem, such as distal sites. Nevertheless, when a complex structure is used, and distal sites are desired, further dissecting the RNA into substructures such as stem loops to carry out several rounds of screening may help solve this problem.

Of all four compounds directly interacted with SL5, Sertraline (H1) showed significant antiviral efficacy against SARS-CoV-2 pseudovirus (PsV) and authentic viral infection in *vitro*, which inhibited SARS-CoV-2 spike (S)-mediated cell-cell fusion through specific binding to the S1 subunit of the viral spike protein, with a particular focus on the receptor-binding domain (RBD) (Chen et al. 2022). Samuraciclib (H2) was ATP-competitive and orally active CDK7 inhibitor, which inhibited the growth of breast cancer cell lines and anti-tumor effects and promoted cell apoptosis (Coombes et al. 2023). There were no studies related to the treatment of the virus as of now. Minocycline (H5) inhibited microbial protein synthesis by binding to the 16S rRNA (Michaelis et al. 2007) and is the only compound reported in this work that was known to interact with RNA directly. Recent research found the precise location of interaction between Minocycline and 16S rRNA could be its double-stranded site (Chukwudi and Good 2016). Minocycline also exhibited effective antiviral activity against some positive-sense RNA viruses in clinical practice, including SARS-CoV-2(Yates et al. 2020; Gironi et al. 2021). Minocycline also inhibited pro-inflammatory cytokines and matrix metalloproteinases that were involved in coronavirus acute infection, therefore may inhibit both the viral replication and the host exuberant inflammatory response (Francini et al. 2020). JG-98 (D6) is an allosteric heat shock protein 70 (Hsp70) inhibitor, which binds to an allosteric site in the NBD to limit ATPase activity and stabilized SBD/client interactions (Rousaki et al. 2011). JG-98 bound to the hepatitis C viral nonstructural protein 5A (NS5A) to block its assembly and proliferation, which played a role in NS5A-augmented internal ribosomal entry site (IRES)-mediated translation of the viral genome (Khachatoorian et al. 2016). It would be very interesting to explore whether these compounds could be applied in treating infection of corona viruses by interfering with RNA function.

Disney’s group also developed ribonuclease targeting chimera (RIBOTAC), comprising an RNA-binder such as a small-molecule drug that selectively bind RNAs conjugated to another small molecule that recruits and locally activates endogenous ribonuclease (RNase) L to induce the enzymatic cleavage of a target RNA (Costales et al. 2018). A small-molecule RIBOTAC had been utilized to degrade the SARS-CoV-2 RNA genome by targeting an attenuator hairpin RNA structure, which is located near the programmed frameshift regulatory element (Haniff et al. 2020a). Recently, coumarin derivatives were recently reported as a warhead in the optimized RIBOTAC design targeting the SL5 motif which robustly degraded SARS-CoV-2 RNA in cellular models at 1 μM and inhibited virus replication at 20 µM in lung epithelial cells (Tang et al. 2023). Similarly, the three small molecules (H1, H2 and H5) discovered in this work could also be modified into RIBOTAC for combating multiple corona viruses.

Up to date, about 30 Fluorescent RNA aptamers have been widely applied in intracellular localization, riboswitches studies, small molecule biosensor design etc. In an example of biosensors, aptamer Spinach was fused to guanine riboswitch through stem-to-stem fusion to detect known ligand, guanine, in the environment as well as in the cell (You et al. 2015). However, this approach has never been applied on detecting novel binders to RNA without previously known ligand. In this work, we took a bold approach believing that RNA-small molecule interaction represents a largely unknown realm, thus deserves more exploration. During the screening, Spinach 2 revealed some disadvantage as a reporter. We found certain compounds could bind to Spinach 2 and displace DFHBI-1T from its binding site, thus reduced fluorescent signal. This may partly arise from the G4 structure in the core of Spinach 2, as G4 has been shown to interact nonspecifically with a number of compounds. Other fluorescent aptamers could be explored to improve this technology in future.

Corona virus genome showed high mutation rates which required constant development of drugs to combat the disease during the pandemic. Although COVID-19 no longer represents a significant threat to human health right now, we could not deny the likelihood that future mutations could lead to new strains jumping from its animal host to human. Hence, developing broad-spectrum drugs for this viral family is necessary. Unlike many existing drug targets, SL5 is not only highly conserved in SARS-CoV-2 genome, but also a conserved feature shared among β corona viruses with several strains infect human. Our work showed the possibility of finding candidates that may serve as broad-spectrum drugs for corona viruses threatening human health in future. In addition, the simple design, robust and fast screening of FAS could assist us in discovering new small molecule drugs targeting all family of RNA viruses and combating other human diseases.

## Materials and methods

### Sequence and structure comparison between SARS-CoV, SARS-CoV-2 and MERS-CoV

Data of SARS-CoV-2 sequences, single nucleotide variation types and frequencies were obtained from the SARS-CoV-2 Mutation Portal, based on 5,340,569 SARS-CoV-2 genome sequences (Saldivar-Espinoza et al. 2023). Genomic RNA sequences of SARS-CoV and MERS-CoV were retrieved from the National Center for Biotechnology Information (NCBI). Mutation types and frequencies of SARS-CoV and MERS-CoV within the apical regions were sourced from previous report (Chen et al. 2021). The secondary structures of SARS-CoV-2, SARS-CoV and MERS were extracted from PDB 8UYS, 8UYP and 8UYK (Kretsch et al. 2024) using RNAPDBee 2.0, respectively. RNA sequence alignment was generate by MEGA and drawn by ENDscript.

### Cloning

The DNA template for RNA synthesis was cloned into the pBSKseph vector, with the template plasmid being the synthesized pBSKseph-Fusion176 (Beijing Liuhe BGI Co., Ltd). Plasmid construction was performed using the KunGre II Multi One Step Cloning Kit (Cat. GT152-02, Greact Biotechnology) for seamless cloning. First, linearized pBSKseph vector was prepared by PCR using Phusion high-fidelity polymerase (Cat. F-5302, Thermo Scientific) with primers Fusion-li F and Fusion-li R (Supplementary table 1), respectively. The product was digested overnight with DpnI (Cat. R0176, NEB) at 37°C to remove the template and then purified by gel extraction (Cat. DC301, Vazyme) to obtain the linearized vector. Next, the insertion fragments were obtained from pBSKseph-Fusion176 plasmid by site-directed mutagenesis PCR using Phusion high-fidelity polymerase and specific primers (Supplementary table 1) and then gel extracted. The ends of insertion fragments bore 15-25 nt sequences homologous to the ends of the linearized vector for homology recombination. The linearized pBSKseph vector and insertion fragments were combined to prepare the 10 µL reaction mixture and incubated at 50°C for 5-15 minutes according to the KunGre II Multi One Step Cloning Kit instructions.

The assembled reaction mixture was then transformed into DH5α competent cells (Cat. CB101, Tiangen) and plated onto LB Agar plates containing 1% ampicillin and incubated at 37°C overnight. Single colonies were selected for sequencing with M13F (Supplementary table 1) universal primer and correct colonies were used to extract plasmids using plasmid mini-prep kit (Cat. DP103, Tiangen) according to the kit instructions.

The synthesized SARS-CoV-2-SL5, MERS-CoV-SL5, SARS-CoV-SL5, DA, DB fragments were cloned into the pBSKseph vector using the same experimental procedure as mentioned above. All primers used in this study were listed in the Supplementary table 1. All plasmids used in this study were listed in the Supplementary table 2.

### In vitro transcription, Purification and refolding of RNA

The DNA template for *in vitro* transcription was first amplified using Phusion high-fidelity polymerase for overhang PCR with specific primers (listed in the Supplementary table 1) to obtain a DNA template containing the T7 promoter. The product was purified using FastPure Gel DNA Extraction Mini Kit. The purified DNA template was sequenced using the IVT-seq primer (Supplementary table 1).

Next, RNA was *in vitro* transcribed using the Quick High Yield RNA Synthesis Kit (Cat. E2050S, NEB) according to the manufacturer’s instructions. The *in vitro* transcription product was then purified using Quick Spin Columns for Radiolabeled RNA Purification (Cat. 11273990001, Merck) following the manufacturer’s instructions. RNA concentration and purity were measured using NanoDrop (Thermo Fisher Scientific, NanoDrop2000).

The purified RNA was diluted in Renature buffer (40 mM HEPES-KOH pH 7.5, 10 mM MgCl_2_, 125 mM KCl) to a final concentration of 3 µM. To ensure consistent folding, the RNA was heated at 75°C for 3 minutes, followed by slow cooling to room temperature.

### Solid phase synthesis, purification and refolding of RNA

For ^19^F-labeled NMR titration experiments, the RNA samples were synthesized using RNA solid-phase synthesizer (SYN-HCY-12P, Qingke) and purified using Glen-Pak columns (60-5100-96, Glen Research) following the manufacturer’s instructions. The resulting RNA samples were exchanged into NMR buffer (40 mM HEPES-KOH pH 7.5, 10 mM MgCl_2_, 125 mM KCl). These samples were refolded by heating at 75°C for 3 minutes, and then slowly cooled to room temperature.

### NMR titration

All NMR experiments were carried out on Bruker Avance 600 MHz spectrometer equipped with 5 mm triple-resonance H/F-TCI cryogenic probe. The samples were measured at 25°C unless otherwise specified. For each titration experiment, the RNA sample was titrated with the small molecule at the indicated molar ratios, and the 1D ^1^H and ^19^F spectra were recorded. All NMR data were processed and analyzed using Topspin. All chemical shifts from the spectra were re-referenced using trifluoroacetic acid (TFA) as the reference. For each titration, all peak intensity from different molar ratios were calibrated by adjusting the peak intensity from TFA to the same scale.

### Optimization of screening conditions

RNA including Fusion 171, Fusion 176 and Spinach 2 were used for determining the screening conditions. Their fluorescence was measured across a range of RNA concentrations from 30 nM to 1 µM and DFHBI-1T concentrations from 1.7 µM to 20 µM, while also assessing the fluorescence at different Mg²⁺ concentrations (0–17.3 mM). Each concentration was diluted in two-fold gradients, using a Versette automated liquid handling workstation (Cat.Versette, Thermo Fisher Scientific). For time dependent folding, 300 nM Spinach 2, Fusion 176, Fusion 177, Fusion 178 and Fusion 179 were incubated in Renature buffer at 25°C and fluorescence was measured every 30 minutes using Spark Multimode Microplate Reader (Cat. Spark, TECAN), with a total of seven measurements (excitation wavelength at 482 nm, the emission wavelength at 505 nm) taken over a 4-hour period, starting at 30 minutes after incubation. Each experiment was repeated three times. Ultimately, after gradient testing, it was determined that the buffer for fluorescence measurement contained: 40 mM HEPES-KCl (pH 7.5), 125 mM KCl, and 10 mM MgCl_2_. In addition, DFHBI-1T (final concentration of 3 µM) and RNA (final concentration at 300 nM) were added for incubation. After all components were added, the samples were mixed and incubated at room temperature in the dark for an appropriate period of time to ensure fluorescence stability. Finally, fluorescence measurement was carried out using Spark Multimode Microplate Reader and the fluorescence intensity was recorded to evaluate the optimal time needed for binding of RNA to DFHBI-1T.

### Screening and compound library

The MCE Bioactive Compound Library (MedChemExpress) was used for screening, and a total of 9,528 compounds were screened. First, RNA (Fusion176 or Spinach 2) and DFHBI-1T was diluted into Renature buffer to the final concentration of 300 nM and 3 µM, respectively. The control group Spinach 2 was incubated for 2 hours, whereas the experimental group Fusion 176 was incubated for 8 hours. The incubation was simultaneously aliquoted into 384-well plates (Cat.781077, Greiner), with 32.5 µL per well using the Versette automated liquid handling system. After the incubation time is up, the first measurement of the fluorescence was carried out using the SPARK multi-functional microplate reader. Then 65 nL of the 10 mM compound stock from the compound library were added into each well using the Echo 650 acoustic pipetting system (Cat.Echo 650, Beckman Coulter), so that the final concentration of the compound was 20 µM. Finally, two hours after adding the compound, the fluorescence was measured again using the microplate reader. The excitation wavelength in fluorescence measurement was set to 482 nm, and the emission wavelength was set to 505 nm, respectively.

### Counter Screening

In counter screening, in addition to the Fusion RNA 176 and Spinach 2 groups, we also set up a control group without RNA to eliminate the influence of the fluorescence of the compounds on the experimental results. The solution system of the RNA-containing group was the same as the overall screening system, while the RNA-free group used DEPC water instead of RNA. The experimental procedures of counter screening were the same as those of screening. The difference was that we used the Echo 650 acoustic pipetting system to add the compounds from the compound library to each well, so that the final concentrations of the compounds were 0.78 µM, 1.56 µM, 3.13 µM, 6.25 µM, 12.5 µM, and 25 µM in sequence.

### Isothermal Calorimetry

The MicroCal PEAQ-ITC instrument (Malvern, PEAQ ITC) was used for ITC analysis. The experiments were conducted at 25℃, with reference power at 10 μcal/s; The feedback was set at High; the stir speed was 750 rpm; the initial delay was 60 seconds and the injection interval was 150 seconds; a total of 19 drops were titrated. Except for the first drop, which had a volume of 0.4μL and a Duration of 0.8 s, each of the remaining 18 drops had a volume of 2 μL and a Duration of 4 seconds. The volume of the sample cell was 280 μL, and the volume of the titration syringe was 60 μL. Each experiment involved titrating the compound RNA sample and the titration buffer and repeated twice. The RNA samples and compounds were added to the cell and syringe, respectively. All compounds were purchased from MedChemExpress Company. For the compound dissolved with DMSO, we first dissolved it into an 80 mM stock concentration with DMSO and then diluted it to the required concentration with Renature buffer. To ensure the consistent buffer condition, RNA was precipitated with ethanol, and then dissolved to 30 µM in Renature buffer containing the same concentration of DMSO. For water-soluble compounds, we dissolved them in DEPC-Treated H_2_O to form a 5 mM stock solution and then diluted them into Renature buffer to reach the target concentration. The RNA preparation process was the same as that for DMSO-soluble compounds, except that RNA was diluted to 30 μM with Renature buffer. Yeast tRNA was purchased (Cat.T8630, Beijing Solarbio Science&Technology Co., Ltd) and dissolved in DEPC-Treated H_2_O to a 4 mM stock concentration, which was then diluted to 250 μM with Renature buffer. In the ITC experiment, the concentrations of the selected compounds and RNA were shown in Supplementary Table 3, respectively. In the data analysis, we subtracted the data of the control group from the titration data of the experimental group and the MicroCal PEAQ-ITC Analysis that came with the instrument Software was used for data analysis (https://www.malvernpanalytical.com/en/products/product-range/microcal-range/microcal-itc-range).

### Data analysis

The code for FAS data processing and analysis in this work was written in Python 3.11 (Rossum 1995) with matplotlib.pyplot used for data visualization (Hunter 2007), numpy for numerical computation (Harris et al. 2020), and pandas for data reading and writing (team 2023). All data processing codes were publicly available on GitHub (https://github.com/HangShiLab/RNA-Data-Analysis). The analysis of IC50 and △Emax was carried out by GraphPad Prism 8 (www.graphpad.com). △ Emax (difference in maximum response value) = 176 (Fusion 176, at 25μmol) – Spi (Spinach 2, at 25μmol), and the IC50 of dose response curve (Ratio of 176 - Ratio of Spi), which was calculated based on Four-Parameter Logistic Model (4PL). Figure 1 was drawn by Biorender (BioRender.com). All secondary structures were generated from the Forna website using the dot-bracket notation (Kerpedjiev et al. 2015) other than that in Figure 6 which was drawn by VARNA (Darty et al. 2009). This paper employed two analytical methods at the initial screen: one was the difference in fluorescence change (i.e., the fluorescence intensity after adding the compound minus the intensity before the addition), and the other was the ratio of the fluorescence change (i.e., the ratio between the fluorescence intensity after the addition to that before the addition). ‘176’ represented the fluorescence of Fusion 176 before administration, ‘176’’ represented the fluorescence of Fusion 176 after administration, ‘Spi’ represented the fluorescence of Spinach 2 before administration, and ‘Spi’’represented the fluorescence of Spinach 2 after administration. Then, the differences were “176’-176” and “Spi’-Spi”, and the ratios were “176’/176” and “Spi’/Spi”, respectively. In our experiment, the standard deviation of the fluorescence range of Spi before dosing was 564.7, the average was 12,003, and the coefficient of variation (CV) was 4.7%. Using this CV as a reference, a cutoff value was established by rounding twice the CV to a convenient integer, in order to delineate the boundaries of fluorescence fluctuation. Therefore, if the fluorescence variation range of Fusion 176 was within 10% (0.90< (176’/176)<1.10), then it was considered that the compound had no effect on Fusion 176. For the compounds that increased the fluorescence of Fusion 176, the selection criterion was that the difference in the increase was greater than approximately two-fold of the CV, that was: (176’/176) > 1.10, and |(176’-176)-(spi’-spi)|/ |(spi’–spi)| > 10%. For the compounds that caused the fluorescence of Fusion 176 to decrease, the selection criteria were: the difference in the decrease was greater than two-fold of the CV (176’/176) < 0.90, and the range from | (176’/176) - (Spi’/Spi) | > 10%.

## Acknowledgements

We thank Dr. Ling Chu and Dr. Chunlai Chen for kind discussions; Dr. Yafeng Kang for initiation assistance at the early exploration of this work; Mrs. Jingjing Rong, Mrs. Tingting Zhang and Dr. Jinfeng He from Center of Pharmaceutical Technology, Dr. Qing Chang, Dr. Yan Liu and Dr. Ning Xu from Technology Center for Protein Sciences, Mr. De Li from Center of Biomedical Analysis, Tsinghua University for kind assistance.

## AUTHOR CONTRIBUTIONS

D, M., X, Y. and S, H. designed the experiments; X, Y., D, M. and Y, W. conducted experiments; X, Y., D, M., X, Y., and S, H. analyzed data; X, Y., D, M., Y, W., X, Y., and S, H. wrote the paper.

## DATA AVAILABILITY

The data on which the conclusions of this article were based are in the article itself. The data analysis codes have been deposited in the GitHub. (https://github.com/HangShiLab/RNA-Data-Analysis)

